# Helios is a marker, not a driver, of human Treg stability

**DOI:** 10.1101/2021.04.27.441702

**Authors:** Avery J. Lam, Prakruti Uday, Jana K. Gillies, Megan K. Levings

**Author notes:** Correspondence, BC Children’s Hospital Research Institute, A4-186, 950 West 28^th^ Ave, Vancouver, BC, V5Z 4H4, Canada.

## Abstract

Regulatory T cell (Treg) therapy holds promise as a potentially curative approach to establish immune tolerance in transplantation and autoimmune disease. An outstanding question is whether therapeutic Tregs have the potential to transdifferentiate into effector T cells and thus exacerbate rather than suppress immune responses. In mice, the transcription factor Helios is thought to promote Tregs lineage stability in a range of inflammatory contexts. In humans, the role of Helios in Tregs is less clear, in part due to the inability to enrich and study subsets of Helios-positive versus Helios-negative Tregs. Using an in vitro expansion system, we found that loss of high Helios expression and emergence of an intermediate Helios (Helios^mid^)-expressing population correlated with Treg destabilization. We then used CRISPR/Cas9 to genetically ablate Helios expression in human naive or memory Tregs and found that Helios-knockout and unedited Tregs were equivalent in their suppressive function and stability in inflammation. Thus, high Helios expression is a marker, but not a driver, of human Treg stability in vitro. These data highlight the importance of monitoring Helios expression in therapeutic Treg manufacturing and provide new insight into the biological function of this transcription factor in human T cells.

## Introduction

CD4^+^FOXP3^+^ regulatory T cells (Tregs) are critical mediators of immune tolerance and homeostasis. Strategies to increase Treg numbers or function by ex vivo expansion for adoptive cell therapy and/or by in situ manipulation with biologics are under investigation in multiple disease contexts (1). In response to microenvironmental cues, Tregs can exhibit functional adaptation or transition into non-suppressive, effector T cells: Treg dysfunction and/or loss of FOXP3 expression has been observed in mouse models of inflammation and human autoimmune disease (2,3). The potential for Tregs to functionally destabilize is a major safety concern for Treg therapy.

The majority (∼70%) of FOXP3^+^ cells in mice and humans express the zinc-finger transcription factor Helios ex vivo (4,5). Based on evidence in mice, this protein is thought to be specifically expressed in thymus-derived Tregs and to support their stable inhibitory activity. Relative to their Helios^−^ counterparts, Helios^+^ Tregs exhibit superior suppressive capacity in vitro (6–8) and in a model of scurfy T cell transfer-mediated colitis (8). Mice with Helios-deficient Tregs develop autoimmune disease at 5–6 months of age, which is accelerated upon immunization, viral inflammation, or lymphopenia (9–12). Helios-deficient Tregs downregulate Foxp3, upregulate the inflammatory cytokines IFN-γ, IL-2, and IL-17 (9–11), and adopt an effector T cell gene signature (12).

In humans, studies of Helios^+^ versus Helios^−^ Tregs have yielded conflicting results. Similar to mouse Tregs, Helios^−^ naive and memory Tregs have higher expression of IFN-γ, IL-2, and IL-17A compared to their Helios^+^ counterparts (4,13–16). Using surrogate markers to enrich for memory Tregs with divergent Helios expression, one study identified Helios-enriched cells as more suppressive ex vivo (14) while another found they were less suppressive (16), compared to their enriched Helios^−^ counterparts. Using single-cell cloning, Helios^+^ naive Tregs were comparably suppressive to Helios^−^ clones (15), but Helios^+^ clones isolated from a mixture of naive and memory Tregs were more suppressive than Helios^−^ clones (16). Epigenetically, relative to Helios^−^ Tregs, Helios^+^ total Tregs exhibited greater demethylation of the Treg-specific demethylation region (TSDR) (13,14), a modification associated with stable FOXP3 expression (17), and loss of Helios expression in naive Tregs correlated with TSDR remethylation (18). On the other hand, another study found that Helios^+^ and Helios^−^ naive Tregs had comparable TSDR demethylation (15). Thus, despite several past studies, it remains unclear to what extent Helios is involved in human Treg stability and function.

Disparities in previous data with human cells may be due in part to the range of indirect methods used to enrich or identify Helios-expressing Tregs. We recently optimized a CRISPR/Cas9-based method to edit human Tregs (19), providing a means to interrogate the function of Helios in lineage-committed naive and memory Tregs using a classical gene knockout-based approach.

## Results and Discussion

### Spontaneous Treg loss of FOXP3 in vitro coincides with the emergence of Helios^mid^ cells

A definitive marker to identify human Tregs is lacking since many Treg-associated proteins, including FOXP3 and Helios, are expressed by activated CD4^+^ T cells (20). However, the kinetics of Helios expression in naive Tregs (nTregs), memory Tregs (mTregs), and conventional CD4^+^CD25^−^ T cells (Tconvs) have not been investigated. Flow-sorted Treg subsets and Tconvs (**Fig. S1A**) were stimulated every 5–7 days following a protocol compatible with efficient CRISPR/Cas9 editing (**Fig. S1B**) (19).

Consistent with their ex vivo expression profiles (13,16), a higher proportion of nTregs expressed Helios compared to mTregs throughout expansion, and the proportion of Helios expression in Tconvs was markedly lower than in nTregs or mTregs (**Fig. 1A–B**) Notably, the proportion of Helios expression declined in all T cell subsets over time (**Fig. 1B**). Consistent with previous reports (20,21), three days post-restimulation, Tconvs exhibited activation-induced upregulation of both FOXP3 and Helios, but only Helios returned to baseline levels (**Fig. 1B**). The transient and overall low expression of Helios in Tconvs relative to Tregs suggests that, while not a definitive marker, Helios may be a more reliable approach than FOXP3 to distinguish Tregs from contaminating Tconvs during in vitro expansion.

**Fig. 1.**
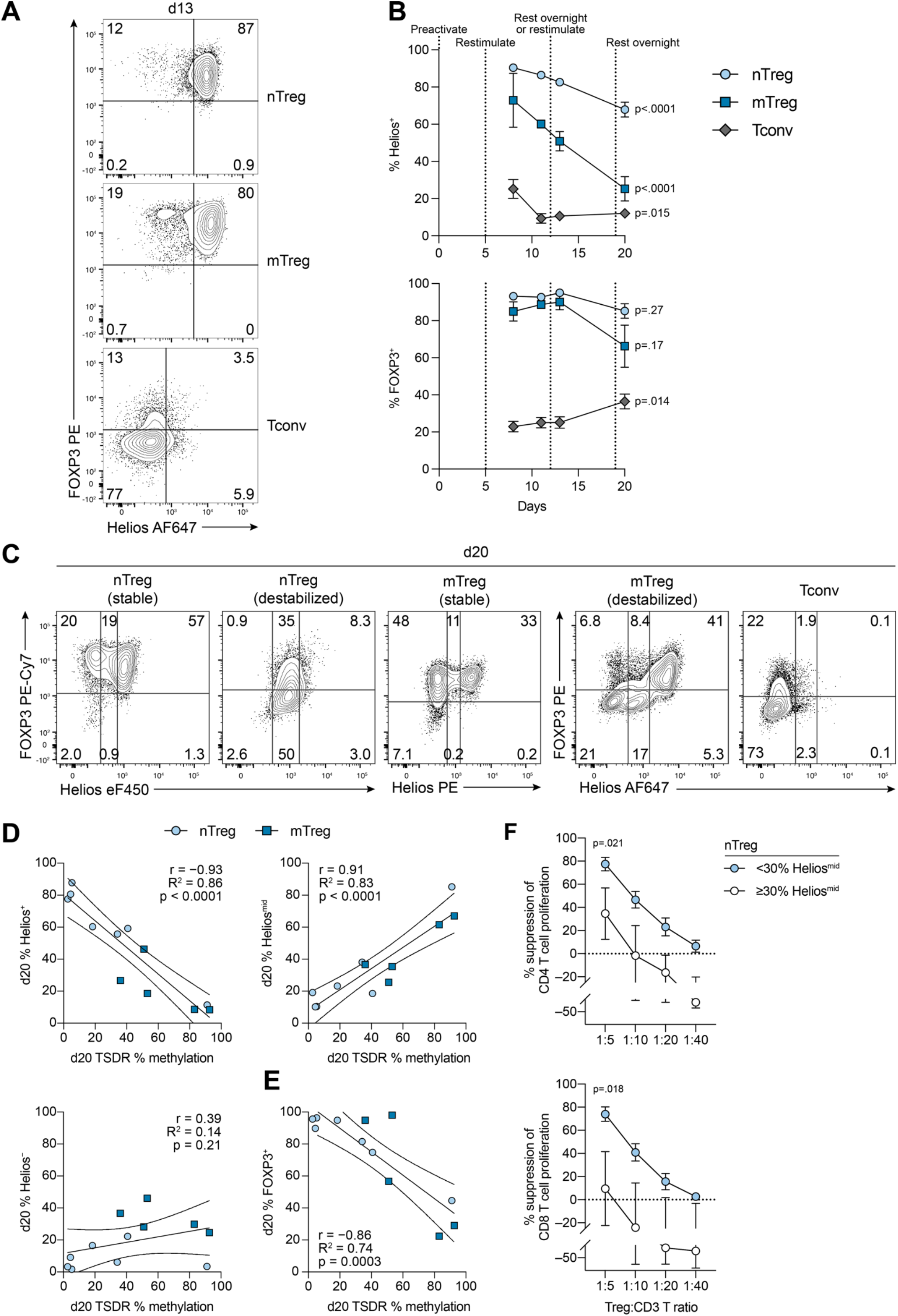
Spontaneous Treg loss of FOXP3 in vitro coincides with the emergence of Helios^mid^ cells. Tregs and Tconvs were preactivated (5 d) and restimulated once or twice with aAPCs (7 d each) before resting overnight. (**A**) Representative Helios and FOXP3 expression at day 13 and (**B**) over time (n=12–20 nTreg, n=2– 14 mTreg, n=11–26 Tconv; 1–16 experiments). (**C**) Representative Helios and FOXP3 expression at day 20 and (**D-E**) correlation with TSDR CpG methylation (day 20) (n=7 nTreg, n=5 mTreg, n=5 experiments). (**F**) Suppression of CD4^+^ and CD8^+^ T cell proliferation by nTregs (day 20) (<30% Helios^mid^, n=6–8; ≥30% Helios^mid^, n=3). (**B, F**) depict mean±SEM. Dots in (**D-E**) represent individual donors and lines represent line of best fit with 95% confidence intervals. Significance determined by 2-way ANOVA with Dunnett’s test comparing day 8 vs. day 20 in (**B**) and t-test of the areas under the curve in (**F**). Pearson correlation coefficient (r) shown in (**D-E**).

We found that occasionally, repetitive TCR activation led to loss of FOXP3 expression in nTregs (n=5 of 20) and mTregs (n=4 of 8), with <80% FOXP3^+^ cells on day 20 (**Fig. 1C**). In addition to loss of FOXP3 protein expression, there was parallel TSDR remethylation and emergence of a population of cells with intermediate Helios expression (Helios^mid^) (**Fig. 1C**). The presence of Helios^mid^ cells correlated strongly with TSDR remethylation, while the proportion of Helios^−^ cells did not (**Fig. 1D**). TSDR remethylation also correlated with loss of FOXP3 expression (**Fig. 1E**). Functionally, nTregs with a high fraction (≥30%) of Helios^mid^-expressing cells were significantly less able to suppress CD4^+^ or CD8^+^ T cell proliferation, compared to nTregs with <30% Helios^mid^ (**Fig. 1F**).

While it is impossible to exclude the possibility that contaminating Tconvs outgrew expanded Tregs, the finding that expanded Tconvs were predominantly Helios^−^ rather than Helios^mid^ (**Fig. 1C**) suggests this is unlikely the case. Furthermore, Tregs that destabilized by day 20 showed high FOXP3 expression and low TSDR methylation on day 13 (**Fig. S2A–B**), and Helios expression at day 13 correlated poorly with subsequent TSDR remethylation (**Fig. S2C**). Thus, features of Treg dysfunction developed during the third round of stimulation, and prior FOXP3 and Helios expression or TSDR methylation does not predict subsequent Treg destabilization.

To our knowledge, intermediate Helios expression (Helios^mid^) in human Tregs has not previously been reported. In mouse colons, subsets of Tregs expressing the transcription factors GATA3 or RORγt can be distinguished by high or low/intermediate expression of Helios, respectively (22); both GATA3^+^ and RORγt^+^ Tregs have distinct functions in gut homeostasis (23). In lymphopenic mice, adoptively transferred Helios^−^ Tregs are more likely to lose Foxp3 expression and can acquire low/intermediate expression of Helios (8), in contrast to adoptively transferred Helios^+^ Tregs which retain both Foxp3 and Helios expression. Whether acquisition of low/intermediate Helios expression and loss of Foxp3 expression are related in this system remains to be determined.

Our observations are in line with the notion that nTregs are less prone to spontaneous FOXP3 loss than mTregs upon repetitive stimulation (**Fig. 1C–D**) (24–28). In previous studies, low mTreg expression of FOXP3 (30–60% FOXP3^+^) in the first 1–2 weeks of expansion (27,28) makes it difficult to rule out the possibility of contaminating Tconvs. Our finding that mTregs sustained high expression of FOXP3 and overall low TSDR methylation for at least two weeks (**Fig. S2A–B**) suggests that Treg impurity is unlikely to be the dominant reason for the observed mTreg instability.

### Helios^KO^ nTregs maintain their phenotype, function, and stability in inflammation

The functional significance of Helios in human Tregs remains controversial. To directly investigate its role in Treg stability, we used CRISPR/Cas9 to knock out Helios in lineage-committed nTregs. As Treg destabilization upon repetitive stimulation was only occasional (25% of nTregs, 50% of mTregs in **Fig. 1B**), we only included Tregs that had not spontaneously lost FOXP3 (≥80% FOXP3^+^ in Cas9 Tregs throughout expansion) for these experiments.

Using two previously validated Helios-targeting gRNAs (29), we found that a combination of both gRNAs yielded the highest Helios protein KO, as assessed three days after editing (**Fig. 2A**). Tregs from mice with a Treg-specific Helios deletion have a survival disadvantage in non-competitive (lymphopenic) and competitive (bone marrow chimera) settings relative to wildtype Tregs, as a result of impaired IL-2-induced STAT5 activation (9) and/or downregulation of the anti-apoptotic protein Bcl-2 (10). In human cells, Helios^KO^ nTregs did not have a competitive disadvantage—the proportions of Helios^+^ and Helios^−^ cells after editing remained the same for two weeks (**Fig. 2B–C**).

**Fig. 2.**
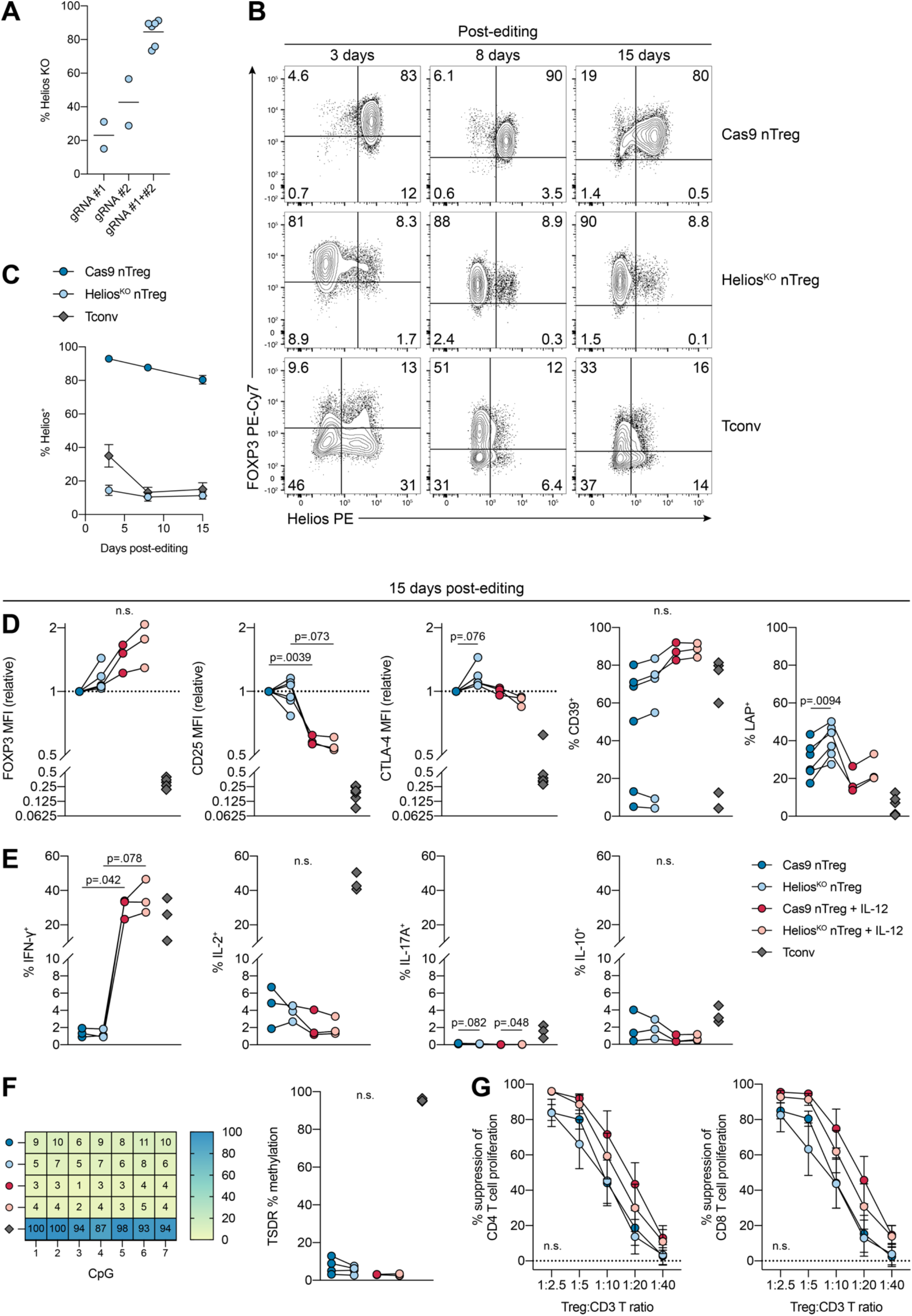
Human Helios^KO^ nTregs maintain their phenotype, function, and lineage stability in inflammation. nTregs were preactivated with aAPCs (5 d), electroporated with Cas9 or Helios-targeting gRNA(s), restimulated twice with aAPCs (7 d each), and rested overnight (total 20 d; n=2–6, 1–3 experiments). (**A**) Extent of Helios protein KO relative to Cas9 nTregs, by flow cytometry. In (**B-G**), gRNAs #1+#2 from (**A**) were used in combination (Helios^KO^). (**B**) Representative Helios and FOXP3 expression and (**C**) average Helios expression over time. In (**D–G**), IL-12 was added every 2–3 d (**Fig. S1B**). (**D**) FOXP3, CD25, CTLA-4, CD39, and LAP expression (day 20). (**E**) IFN-γ, IL-2, IL-17A, and IL-10 expression (day 20) after restimulation with PMA, ionomycin, and brefeldin A (6 h). (**F**) Representative and average TSDR CpG methylation. (**G**) Suppression of CD4^+^ and CD8^+^ T cell proliferation by Tregs (day 20). Dots in (**A, D-F**) depict individual donors. (**C, G**) depict mean±SEM. Significance determined by mixed-effects model with Geisser-Greenhouse correction and Sidak’s test in (**D-F**) and one-way ANOVA of the areas under the curve with Sidak’s test in (**G**). Tconvs shown in (**D-F**) for reference.

We reasoned that phenotypic and functional consequences of Helios^KO^ may arise in a progressive manner and/or under inflammatory conditions. Mice with Helios-deficient Tregs develop autoimmunity characterized by excessive Th1 and B cell responses, but with delayed kinetics relative to Treg-deficient mice (9,10,30). Thus, we expanded Tregs for two weeks after Helios^KO^ in the presence of the inflammatory cytokine IL-12, which is known to cause mouse and human Tregs to adopt a Th1-like, IFN-γ-producing phenotype (31–36), with some studies reporting reduced suppressive function (32,34,36).

Human Helios^KO^ nTregs retained a Treg phenotype in Th1 inflammation. Helios^KO^ nTregs transiently upregulated FOXP3 (**Fig. S3A**) but this difference was not significant after two weeks (**Fig. 2D**). IL-12 exposure decreased CD25 expression but in a similar manner in Cas9 and Helios^KO^ nTregs (**Fig. 2D, Fig. S3A**). Additionally, we found no differences in viability between cell types with or without IL-12 (data not shown). Consistent with the activated phenotype of mouse Helios-deficient Tregs (10,12), we found sustained upregulation of the latent TGF-β-associated molecule LAP upon Helios^KO^ in nTregs (**Fig. 2D, Fig. S3A**). LAP marks a subset of highly suppressive Tregs (37). Other Treg functional proteins including CTLA-4 and CD39 were unaffected by either Helios^KO^ or IL-12 exposure (**Fig. 2D, Fig. S3A**).

As expected, the addition of IL-12 substantially increased nTreg expression of IFN-γ, particularly after two weeks (**Fig. 2E, Fig. S3B**), but to a similar extent in both Cas9 and Helios^KO^ cells. Expression of other cytokines, including IL-2, IL-17A, and IL-10, remained low (**Fig. 2E, Fig. S3B**), despite previous evidence that Helios transcriptionally represses *Il2* expression in mouse Tregs (38). These observations in human cells differ from those with mouse Helios-deficient Tregs, which not only downregulated Foxp3 but also displayed elevated IFN-γ, IL-2, and IL-17 expression compared to wildtype Tregs (9–12).

Because loss of high Helios expression correlated with TSDR remethylation in nTregs and mTregs, either upon repetitive in vitro stimulation (**Fig. 1D**) or induced by 4-1BB or TNFR2 signals (18), we asked whether Helios^KO^ affected TSDR methylation. Helios^KO^ nTregs maintained a highly demethylated TSDR similar to Cas9 nTregs, even with prolonged exposure to IL-12 (**Fig. 2F, Fig. S3D**). These data are in line with unchanged or increased FOXP3 protein expression (**Fig. 2D, Fig. S3A**).

Functionally, Cas9 and Helios^KO^ nTregs exhibited equivalent, high suppressive capacity of CD4^+^ and CD8^+^ T cell proliferation in vitro (**Fig. 2G, Fig. S3D**). IL-12 exposure increased Treg suppression of CD4^+^ T cells (**Fig. S3D**), while prolonged IL-12 exposure (two weeks) revealed no significant differences in suppressive function (**Fig. 2G**).

In mice, Helios-deficient Tregs downregulate Foxp3 expression (9,10), but one group found no accompanying change in TSDR demethylation (10), suggesting that other mechanisms may be responsible for the observed Foxp3 loss in that model. In human Tregs, two previous studies used siRNA to knockdown Helios with varying efficiencies and effects: one group reported no change in suppressive function (4), while another found reduced function with Helios knockdown (5). As siRNA was delivered by electroporation and hence had only transient gene repression, Treg function could only be assessed within 24–60 h. Here, we used CRISPR/Cas9 to stably ablate Helios for up to two weeks (**Fig. 2B–C**), revealing that Helios expression is not required for human Treg suppressive function, at least as assessed by suppression of T cell proliferation in vitro.

### Human Helios^KO^ mTregs retain a Treg phenotype

As mTregs are less stable in vitro and show a more heterogenous Helios expression pattern than nTregs (**Fig. 1D–E**) (13,16), we asked whether Helios has a distinct role in mTregs. We expanded mTregs for two weeks, due to their propensity to spontaneously destabilize by the third week even without any genetic perturbation (**Fig. 1D–E**). Using CRISPR/Cas9, we were able to achieve a similar protein KO efficiency (**Fig. 3A–B**) and reduction in Helios expression in mTregs (**Fig. 3C**) as in nTregs. Of note, since mTregs comprise a mixture of Helios^+^ and Helios^−^ cells, mTregs after editing are a mixture of endogenous Helios^−^ cells and Helios^KO^ mTregs, and the two populations may not be functionally equivalent (8).

**Fig. 3.**
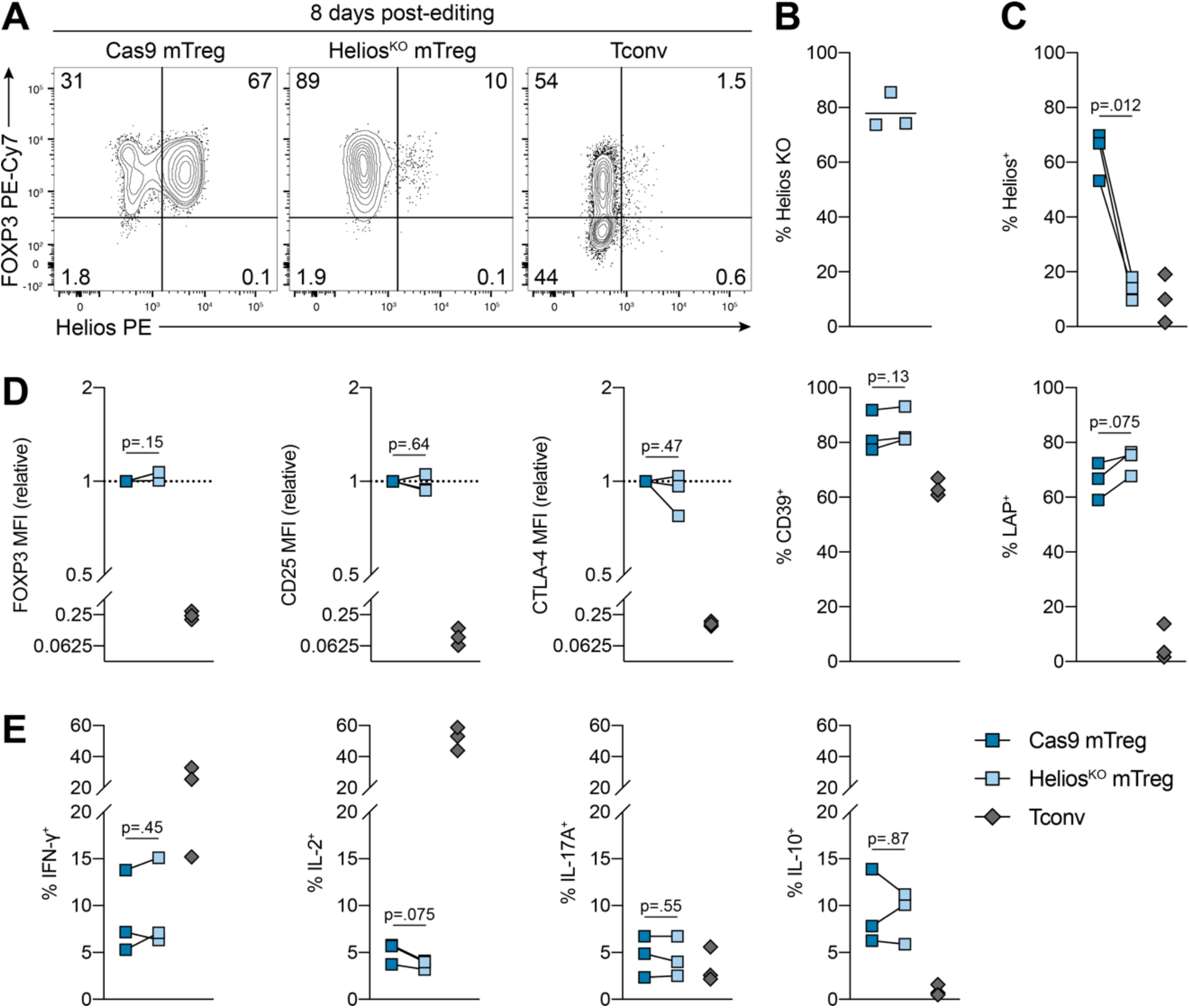
Human Helios^KO^ mTregs retain a Treg phenotype. mTregs were preactivated with aAPCs (5 d), electroporated with Cas9 or Helios-targeting gRNAs #1+#2 (Helios^KO^), restimulated with aAPCs (7 d), then rested overnight (total 13 d, n=3, 1 experiment). (**A**) Representative Helios and FOXP3 expression. (**B**) Amount of Helios protein KO relative to Cas9 mTregs. (**C**) Helios expression. (**D**) FOXP3, CD25, CTLA-4, CD39, and LAP expression. (**E**) IFN-γ, IL-2, IL-17A, and IL-10 expression after restimulation with PMA, ionomycin, and brefeldin A (6 h). Dots in (**B-E**) represent individual donors. Significance in (**C-E**) determined by paired t-test. Tconvs shown for reference.

Helios^KO^ mTregs maintained expression of FOXP3 and CD25, as well as the functional proteins CTLA-4, CD39, and LAP, compared to Cas9 mTregs (**Fig. 3D**). Analogous to nTregs, Helios^KO^ mTregs exhibited a trend towards LAP upregulation, though this difference was not significant (**Fig. 3D**). We then investigated mTreg expression of effector cytokines. Although human Helios^−^ Tregs, which are enriched for mTregs, have been reported to upregulate low levels of IFN-γ, IL-2, and IL-17A (14,16), we did not find a further increase in their expression upon Helios^KO^, suggesting that Helios is uncoupled from cytokine downregulation in mTregs. Overall, Helios ablation in human mTregs did not affect their expected Treg phenotype.

### Concluding Remarks

In summary, Helios identifies, but is not required for, stable human Tregs. Specifically, we found: (i) stimulation of human Tregs three times can lead to loss of FOXP3 expression in vitro (25% of nTreg donors, 50% of mTreg donors); (ii) emergence of a Helios^mid^ population in nTregs and mTregs correlates with their phenotypic, epigenetic, and functional destabilization; (iii) nevertheless, Helios ablation does not negatively affect nTreg or mTreg phenotype, function, or stability in inflammation. In the context of Treg cell therapy, it may be instructive to include Helios in addition to FOXP3 as a marker during Treg expansion to monitor their stability and purity, particularly given the transient nature of activation-induced upregulation of Helios in Tconvs. Strategies to target or stabilize Helios in mature Tregs, however, may not be as critical.

One major difference from studies in mice is that Helios is ablated during mouse Treg development (9–12), whereas in our human study, Helios was ablated in lineage-committed Tregs. Rather than serving an ongoing function in Tregs, Helios may instead be important for establishing Treg lineage commitment. This possibility is supported by the finding that human fetal naive T cells deficient in Helios are unable to differentiate into bona fide Tregs (29). Similarly, our investigation into the role of FOXP3 in mature human Tregs revealed that continuous expression of this master transcription factor is dispensable for maintenance of human Treg identity (19). Collectively, our study and those of others suggest that the dominant role for Helios is during human Treg development, and that although its expression marks stable Tregs, it does not have a direct role in maintaining this lineage-committed state.

### Statements

The data that support the findings of this study are available from the corresponding author upon reasonable request. MKL has received research funding from Sangamo Therapeutics, Bristol-Myers Squibb, Pfizer, Takeda, and CRISPR Therapeutics for work unrelated to this study. All other authors declare no competing interests. Collection of human samples was performed after written informed consent in accordance to protocols approved by the University of British Columbia Clinical Research Ethics Board and Canadian Blood Services Research Ethics Board.

## Author Contributions

Conceptualization: AJL, MKL

Formal analysis: AJL, JG

Funding acquisition: MKL

Investigation: AJL, PU, JG

Supervision: MKL

Writing—Original Draft: AJL

Writing—Review and Editing: AJL, PU, JKG, MKL

## Acknowledgements

We thank BC Children’s Hospital Research Institute (BCCHR) Flow Core Facility for assistance with flow sorting. This work was supported by a grant from the Canadian Institutes of Health Research (CIHR; FDN-154304 to MKL). AJL is supported by a CIHR Doctoral Research Award and MKL receives a BCCHR salary award.

## Materials and Methods

### CRISPR/Cas9 editing reagents

*IKZF2* (Helios)-targeting gRNAs have been previously described (29). gRNA target sequences: IKZF2 gRNA #1, 5’-GGAGGAATCCGGCTTCCGAA-3’; IKZF2 gRNA #2, 5’-GATACTACCAAGGCTCCTAA-3’. gRNA (100 μM) was generated by duplexing crRNA and tracrRNA (both IDT) at a 1:1 molar ratio for 5 min at 95°C, followed by gradual cooling to room temperature. CRISPR/Cas9 ribonucleoprotein was generated by combining gRNA and SpCas9-2xNLS (40 μM; QB3 MacroLab) at a 2:1 molar ratio for 10 min at room temperature.

### T cell isolation

Cells were sequentially enriched for CD4^+^ and CD25^+^ cells by RosetteSep (STEMCELL Technologies) and CD25 MicroBeads II (Miltenyi Biotec), respectively. Naive and memory Tregs were flow-sorted from CD25-enriched cells with a MoFlo Astrios (Beckman Coulter) or FACSAria IIu (BD Biosciences). Tconvs were flow-sorted from the CD25-depleted population. Sorting strategy (**Fig. S1A**): nTregs (CD4^+^CD25^hi^CD45RA^-^CD127^lo^), mTregs (CD4^+^CD25^hi^CD45RA^-^CD127^lo^), Tconvs (CD4^+^CD25^lo^CD127^hi^). CD3^+^ T cells were isolated by RosetteSep (STEMCELL Technologies).

### Treg expansion with CRISPR editing

All cells were cultured in X-VIVO 15 (Lonza) supplemented with 5% (v/v) HS (WISENT), 1% (v/v) penicillin-streptomycin (Gibco), 2 mM (v/v) GlutaMAX (Gibco), 15.97 mg/L phenol red (Sigma-Aldrich) at 37°C, 5% CO_2_. During expansion, media was additionally supplemented with IL-2 (Proleukin; 1000 IU/ml for Tregs, 100 IU/ml for Tconvs) and refreshed every 2–3 days.

Treg expansion with CRISPR editing was performed as described previously (**Fig. S1B**) (19). Briefly, cells were preactivated for 5 days with IL-2 and either 1:1 aAPCs or 25 μl/ml CD3/CD28/CD2 tetramer (ImmunoCult Human CD3/CD28/CD2 T Cell Activator, STEMCELL Technologies). L cells (ATCC CRL-2648) overexpressing CD32, CD80, and CD58, gamma-irradiated (75 Gy), and loaded with anti-CD3 (OKT3, 0.1 μg/ml; University of British Columbia Antibody Lab) served as aAPCs (39).

CRISPR/Cas9 reagents were delivered by electroporation with a Neon Transfection System 10 μL Kit (Invitrogen). Tregs were washed twice with PBS, resuspended in Buffer T (≤20×10^6^ cells/ml), electroporated at 1400 V / 30 ms / 1 pulse with Cas9 (20 pmol per transfection) or Helios-targeting gRNAs (40 pmol gRNA + 20 pmol Cas9, per unique ribonucleoprotein per transfection). Electroporated cells were immediately transferred into prewarmed antibiotic-free media and expanded for 7 days with IL-2 and aAPCs. Preactivated Tconvs were expanded in parallel without electroporation.

12 day-expanded cells were either rested overnight with reduced IL-2 (100 IU/ml for Tregs, none for Tconvs) for phenotypic and functional assays (total 13 days) or further expanded for another 7 days with IL-2 and aAPCs as above before resting overnight (total 20 days). In some cases, recombinant human IL-12 (10 ng/ml, BD Biosciences) was added on days 5, 8, 10, 12, 15, and 17, as indicated (**Fig. S1B**).

nTregs and mTregs in **Fig. 1, Fig. S1D**, and **Fig. S2** were transfected with Cas9. During optimization of Treg expansion with CRISPR/Cas9 editing, we found superior Treg expansion with aAPC-based preactivation relative to CD3/CD28/CD2 tetramers; in both scenarios, cells were restimulated once or twice with aAPCs as above (19). Comparing the two preactivation methods, we found that aAPC-preactivated nTregs, mTregs, and Tconvs displayed similar FOXP3 and Helios kinetics (not shown), and nTregs exhibited comparable suppressive function at day 20 in vitro (**Fig. S1A–B**). As the two methods were comparable, we pooled data from tetramer- and aAPC-preactivated Treg expansions in **Fig. 1** and **Fig. S2**. For experiments investigating the effects of Helios^KO^ on nTregs and mTregs, we used aAPC-based preactivation and restimulation (**Fig. 2, Fig. 3, Fig. S3**).

### In vitro suppression of T cell proliferation

Allogeneic CD3^+^ T cells (responder cells) and expanded Tregs were labelled with Cell Proliferation Dye eF450 and eF670 (Invitrogen), respectively, then cocultured at the indicated ratios and activated with anti-CD3/CD28-coated Dynabeads (Gibco Dynabeads Human T-Expander, 1 bead:16 CD3^+^ T cells) for 4 days. CD3^+^ T cells activated alone served as positive controls. In all cases, CD3^+^ T cells were plated at 50,000 cells/well in a 96-well U-bottom plate. IL-12 was not added during cocultures. Percent suppression of CD4^+^ and CD8^+^ T cell proliferation (gating strategy in **Fig. S1C**) was calculated using division index: (1 – (division index of sample / division index of positive control)) * 100%.

### Flow cytometry

Antibodies are listed in **Table S1**. Cells were stained for surface antigens in PBS (Gibco) or Brilliant Stain Buffer (BD Biosciences) for 20 min at room temperature; Fixable Viability Dye eF780 was used to exclude dead cells. Cells were fixed and permeabilized with the eBioscience Foxp3 / Transcription Factor Staining Buffer Set (Invitrogen) for 40 min at room temperature, washed twice, then stained for intracellular antigens for 40 min at room temperature. For cytokine expression, cells were restimulated with PMA (10 ng/ml), ionomycin (500 μg/ml), and brefeldin A (10 μg/ml; all Sigma-Aldrich) for 6 h at 37°C before staining. Samples were acquired with a FACSymphony A5, LSRFortessa X-20 (both BD Biosciences), or CytoFLEX (Beckman Coulter), and data analysed by FlowJo software (v10.7.1; BD).

All events were pre-gated on live single CD4^+^ cells (**Fig. S1A**), except for assays determining suppression of T cell proliferation (gating strategy in **Fig. S1C**). FOXP3^+^ gates were set based on Tconvs within each donor. Helios^+^ gates were set based on nTregs (for nTregs and mTregs) and Tconvs (for Tconvs) within each donor, and Helios^mid^ cells were defined as the population between these two thresholds. Percent Helios protein KO was calculated as: (1 – (% Helios^+^ of edited sample / % Helios^+^ of unedited sample)) * 100%.

### TSDR methylation

Genomic DNA was isolated from Treg and Tconv samples from male donors, bisulfite-converted with a EZ DNA Methylation-Direct Kit (Zymo Research), and PCR-amplified. Pyrosequencing was performed with a PyroMark Q96 MD (QIAGEN). Primer sequences: forward, 5’-AGAAATTTGTGGGGTGGGGTAT-3’; reverse, 5’-ATCTACATCTAAACCCTATTATCACAACC-3’ (biotinylated); pyrosequencing, 5’-AGAAATTTGTGGGGTGGG-3’.

### Statistical analysis

Normality was assumed. Statistical significance was determined by a paired t-test, mixed-effects model with Geisser-Greenhouse correction and Sidak’s multiple-comparisons test, or 2-way ANOVA with Dunnett’s multiple-comparisons test, as applicable. For Treg-mediated suppression of T cell proliferation, areas under the curve were compared with a t-test or one-way ANOVA with Sidak’s multiple-comparisons test, taking into account all degrees of freedom (40). p < 0.05 was considered significant. Analysis was performed with Prism v9.0.1.

## Abbreviations

aAPC: artificial antigen-presenting cell
mTreg: memory regulatory T cell
nTreg: naive regulatory T cell
Tconv: CD4^+^ conventional T cell
Treg: regulatory T cell
TSDR: Treg-specific demethylation region

