## Supplemental Figures & Tables for "Helios is a marker, not a driver, of human Treg stability"

**Supporting Information for Lam et al., "Helios is a marker, not a driver, of human Treg stability"**

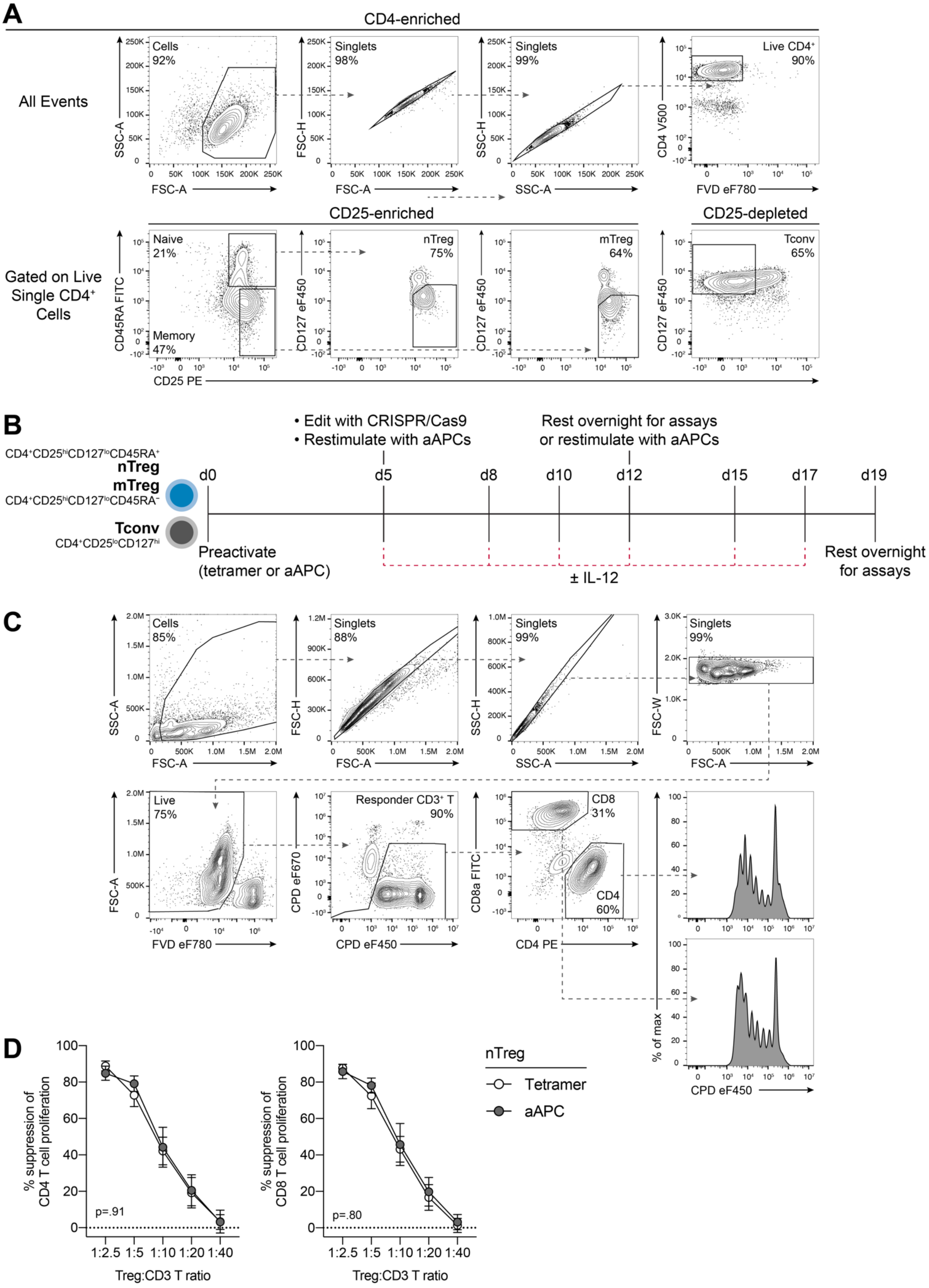

***Fig. S1. Strategies for flow cytometric sorting, gating, and Treg expansion.***

(**A**) Flow sorting strategy for nTregs ($CD4^{+}CD25^{hi}CD45RA^{+}CD127^{lo}$), mTregs ($CD4^{+}CD25^{hi}CD45RA^{-}CD127^{lo}$; both from CD4- and CD25-enriched PBMCs) and Tconvs (from CD4-enriched, CD25-depleted PBMCs). (**B**) Treg and Tconv expansion timeline. Flow-sorted cells were preactivated with CD3/CD28/CD2 tetramer or aAPCs (5 d), edited with CRISPR/Cas9, and restimulated with aAPCs (7 d). Cells were either rested overnight with reduced IL-2 (for day 13) or restimulated again with aAPCs (7 d) before overnight rest (for day 20). In some cases, IL-12 (10 ng/ml) was added every 2–3 days after editing. In all cases, IL-2 (1000 IU/ml for Tregs, 100 IU/ml for Tconvs) was added every 2–3 days throughout expansion except during overnight rest (100 IU/ml for Tregs, none for Tconvs). (**C**) Gating strategy to evaluate Treg-mediated suppression of $CD4^{+}$ and $CD8^{+}$ T cell proliferation after 4-day coculture of Tregs (CPD eF670-labelled) and allogeneic $CD3^{+}$ T cells (CPD eF450-labelled) with anti-CD3/CD28-coated Dynabeads. (**D**) nTregs were preactivated with CD3/CD28/CD2 tetramers or aAPCs as indicated, then restimulated twice with aAPCs (7 d each) before overnight rest (for day 20). Suppression of $CD4^{+}$ and $CD8^{+}$ T cell proliferation (n=6–8 Tetramer, n=5–7 aAPC, 3–4 experiments). Significance in (**D**) determined by t-test of the areas under the curve. nTreg, naive Treg; mTreg, memory Treg; Tconv, conventional T cell; FVD, Fixable Viability Dye; CPD, Cell Proliferation Dye.

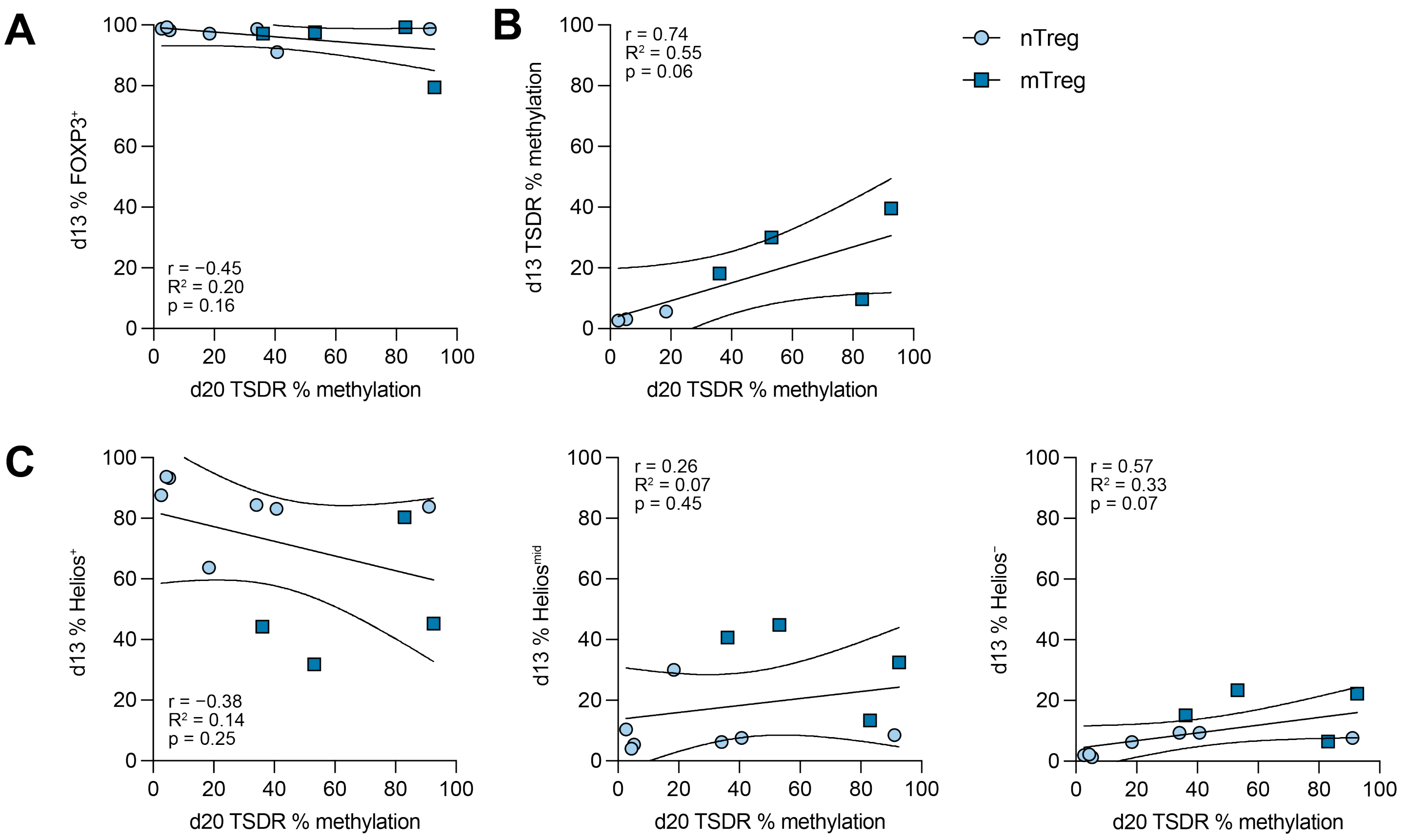


***Fig. S2. Association of Treg instability with suppressive function, prior Helios and FOXP3 expression, and prior TSDR methylation.***

Flow-sorted nTregs and mTregs were preactivated (5 d), restimulated with aAPCs (7 d), and either rested overnight (for day 13) or restimulated again with aAPCs (7 d) before overnight rest (for day 20) (n=3–7 nTreg, n=4 mTreg, 5 experiments). (**A-C**) Correlation of TSDR methylation (average of 7 CpGs) at day 20 with (**A**) FOXP3 expression at day 13, (**B**) TSDR methylation at day 13, and (**C**) Helios expression at day 13. Pearson correlation coefficient (r) shown in (**A-C**).

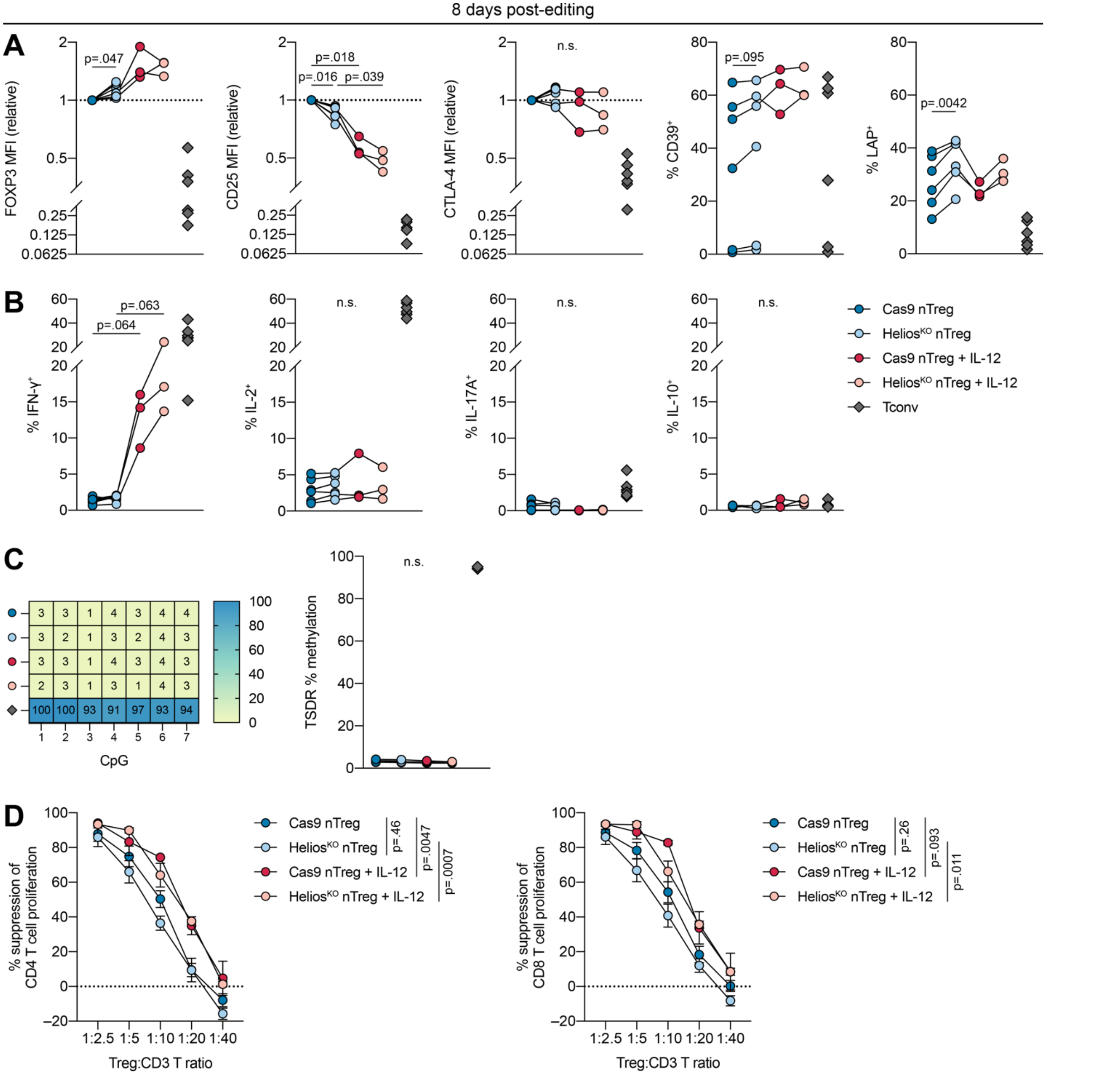


***Fig. S3. Effects of Helios$^{KO}$ on nTreg phenotype, function, and lineage stability in inflammation, 8 days after editing.***

Flow-sorted nTregs were preactivated with aAPCs (5 d), edited with Cas9 or Helios-targeting gRNAs #1+#2 (Helios$^{KO}$), restimulated with aAPCs (7 d), and rested overnight with reduced IL-2 (n=3–6, 1–2 experiments). Tconvs were expanded in parallel without electroporation. IL-12 was added every 2–3 days (**Fig. S1B**). (**A**) FOXP3, CD25, CTLA-4, CD39, LAP expression. (**B**) IFN-γ, IL-2, IL-17A, IL-10 expression after restimulation with PMA, ionomycin, and brefeldin A (6 h). (**C**) Example and average TSDR methylation of 7 CpGs. (**D**) Suppression of CD4$^+$ and CD8$^+$ T cell proliferation. Dots in (**A-C**) depict individual donors. (**D**) depicts mean±SEM. Significance determined by mixed-effects model with Geisser-Greenhouse correction and Sidak's test in (**A-C**) and one-way ANOVA of the areas under the curve with Sidak's test in (**D**). Tconvs shown in (**A-C**) for reference.

***Table S1. Antibodies used for flow cytometry.***

| Antigen | Clone | Fluorophore | Company |
|---|---|---|---|
| CD4 | OKT4 | PE | Invitrogen |
| CD4 | RPA-T4 | BV605 | BD Biosciences |
| CD4 | RPA-T4 | FITC | Invitrogen |
| CD4 | RPA-T4 | V500 | BD Biosciences |
| CD4 | SK3 | BUV395 | BD Biosciences |
| CD4 | SK3 | BUV496 | BD Biosciences |
| CD25 | 4E3 | PE | Miltenyi Biotec |
| CD25 | 2A3 | BUV395 | BD Biosciences |
| CD25 | 2A3 | BV711 | BD Biosciences |
| CD39 | A1 | BV510 | BioLegend |
| CD39 | A1 | FITC | BioLegend |
| CD8a | HIT8a | FITC | Invitrogen |
| CD45RA | HI100 | FITC | BD Biosciences |
| CD127 | eBioRDR5 | eF450 | Invitrogen |
| CD152 (CTLA-4) | BNI3 | APC | BD Biosciences |
| CD152 (CTLA-4) | BNI3 | BV786 | BD Biosciences |
| FOXP3 | 236A/E7 | PE | Invitrogen |
| FOXP3 | 236A/E7 | PE-Cy7 | Invitrogen |
| Helios | 22F6 | AF488 | BioLegend |
| Helios | 22F6 | AF647 | BioLegend |
| Helios | 22F6 | eF450 | BioLegend |
| Helios | 22F6 | PE | BioLegend |
| IFN-γ | B27 | BUV395 | BD Biosciences |
| IL-2 | MQ1-17H12 | BUV737 | BD Biosciences |
| IL-10 | JES3-9D7 | BV421 | BioLegend |
| IL-17A | N49-653 | BV786 | BD Biosciences |
| LAP | FNLAP | PE-Cy7 | Invitrogen |
